# Astrocyte reprogramming drives tumor progression and chemotherapy resistance in agent-based models of breast cancer brain metastases

**DOI:** 10.1101/2025.06.05.654750

**Authors:** Rupleen Kaur, Rowan Barker-Clarke, Andrew Dhawan

## Abstract

Breast cancer brain metastases (BCBM) affect nearly 90,000 patients annually in the United States and carry a significant risk of mortality. As metastatic lesions develop, the unique milieu of the brain microenvironment shapes disease progression and therapeutic response. Among resident brain cells, astrocytes are both the most common, and are increasingly recognized as key regulators of this process, yet their precise role remains poorly defined. Here, we present a hybrid agent-based model (ABM) to simulate tumor–astrocyte interactions on a two-dimensional lattice. In our model, metastatic tumor cells induce phenotypic reprogramming of astrocytes from an anti-to a pro-metastatic state, thereby enhancing tumor proliferation. We systematically evaluate how variations in astrocyte density, spatial distribution, and chemotherapy impact tumor expansion and spatial morphology, quantified by fractal dimension, lacunarity, and eccentricity. Our simulations reveal that astrocyte reprogramming accelerates tumor progression and contributes to increased morphological complexity and chemotherapeutic resistance.

## Introduction

Breast cancer brain metastases (BCBM) affect approximately 20- 30% of patients with metastatic breast cancer and severely impact both survival and quality of life Riecke et al. (2023). Patients with BCBM have a median overall survival of 14 months Shen et al. (2015). Despite advances in systemic therapy, BCBM are difficult to treat, in part due to the unique physiological and cellular characteristics of the brain microenvironment Wang et al. (2021).

The BCBM microenvironment is comprised of glial cells (astrocytes, oligodendrocytes, and microglia) and neurons Ishibashi and Hirata (2024). Astrocytes, the predominant glial cell type in the brain, have emerged as pivotal modulators of tumor progression Kaverina et al. (2017); Kaur et al. (2025). When tumor is not present, astrocytes maintain neural homeostasis and support blood–brain barrier integrity Liu et al. (2024); Valles et al. (2023). In the context of BCBM, astrocytes initially adopt an anti-metastatic phenotype, restricting tumor outgrowth by secreting plasminogen activators that promote apoptosis in disseminated cancer cells that have not yet adapted to the brain microenvironment Valiente et al. (2014). However, some cancer cells develop resistance by upregulating plasminogen activator inhibitors, including SerpinB2 and neuroserpin, thereby evading astrocyte-mediated cytotoxicity Valiente et al. (2014). Subsequently, tumor-derived signals reprogram astrocytes into a pro-metastatic state Burn et al. (2021); Klein et al. (2015); Wasilewski et al. (2017). This pro-metastatic state results in increased tumor proliferation via the secretion of interleukins (IL-6, IL1*β*), tumor necrosis factor-*α* (TNF-*α*), and fatty acids, which activate STAT3, NF-*κ*B, and PPAR*γ* signaling pathways both in co-culture and in *in vivo* models of BCBM Zou et al. (2019); Seike et al. (2010); Wasilewski et al. (2017).

Tumor cells also establish gap junctions with astrocytes — primarily via connexin 43 — which facilitates the cell-cell transfer of signaling molecules such as cGAMP and reduces intracellular calcium, thereby promoting survival and resistance to chemotherapy-induced apoptosis Chen et al. (2016); Lin et al. (2010). This suggests that astrocyte-tumor gap junction signaling may contribute to the emergence of localized ‘chemo-protective pockets’ within BCBM lesions, where gap junction–mediated sequestration of pro-apoptotic signals—including calcium—could spatially restrict drug efficacy and promote tumor cell survival under chemotherapeutic stress. While the molecular components of these interactions are increasingly well-characterized Seike et al. (2010); Wasilewski et al. (2017), a spatially resolved understanding of how astrocyte–tumor crosstalk regulates tumor dynamics and treatment response in BCBM remains lacking. Such spatial insights are critical for identifying targetable niches of therapeutic resistance within the brain microenvironment.

The brain is a highly heterogeneous organ, both anatomically and at the cellular level Lee et al. (2022). Astrocyte density and morphology differ markedly across brain regions: gray matter contains densely packed protoplasmic astrocytes with complex arborization, while white matter is populated by fibrous astrocytes with distinct structural and functional properties Köhler et al. (2019); Bocchi et al. (2025). More-over, regional variability in astrocyte density has been reported across multiple brain regions, including the cortex, hippocampus, cerebellum, and brainstem, each of which could therefore provide a distinct metastatic niche for invading tumor cells Man et al. (2024); Endo et al. (2022); Forrest et al. (2022). These spatial differences may influence how tumor cells interact with resident astrocytes and respond to therapy. However, these region-specific dynamics remain difficult to resolve using traditional experimental approaches alone, underscoring the need for computational frameworks capable of cintegrating spatial heterogeneity and capturing emergent behaviors in tumor–astrocyte ecosystems.

Here, we develop a two-dimensional, hybrid agent-based model (ABM) to simulate tumor–astrocyte interactions within the BCBM microenvironment. While previous ABMs modeled tumor growth, invasion, and treatment response in other tissue contexts, prior ABMs have not been applied to examine astrocyte behavior in brain metastases Robertson-Tessi et al. (2015); Shyntar et al. (2022); West et al. (2024). We model tumor cells and astrocytes on a two-dimensional lattice as autonomous agents governed by biologically informed rules, accounting for tumor cell proliferation, treatment effects, and tumor-astrocyte crosstalk. This framework allows for the systematic evaluation of how astrocyte density and spatial distribution influence tumor expansion, spatial morphology, and treatment resistance.

Using this model, we demonstrate that astrocyte reprogramming may be a key driver of BCBM progression, and can affect treatment response. By varying astrocyte density, spatial distribution, and functional state, we show the extents to which microenvironmental conditions can influence BCBM growth, spatial structure, and sensitivity to therapy. Within the framework of our model, astrocyte reprogramming accelerates tumor expansion, increases morphological complexity, and promotes spatial configurations associated with aggressive behavior. Moreover, we show that astrocyte–tumor interactions reduce chemotherapy efficacy, suggesting that targeting astrocyte reprogramming may be a reasonable focus for future therapeutics in BCBM.

## Results

### Astrocyte Reprogramming Accelerates Tumor Growth and Alters Spatial Morphology

To assess the effect of astrocyte reprogramming on tumor progression, we conducted agent-based simulations under a fixed parameter set representative of a pro-metastatic regime, comparing conditions with and without astrocyte state switching (Fig. 2A-C). In the absence of state switching, tumors exhibited moderate growth, reaching a mean final size of 8980 ± 576 cells over 50 replicates. In contrast, allowing for astrocyte reprogramming led to a substantial increase in tumor burden, with tumors growing to a mean final size of 20934 ± 939 cells, representing a 133% increase compared to the non-reprogrammable condition (Mann–Whitney *U* = 0.0, *p* = 7.06 *×* 10^*−*18^). Group comparisons were performed using the Mann–Whitney U test to avoid assumptions of normality and equal variance.

**Fig. 1.**
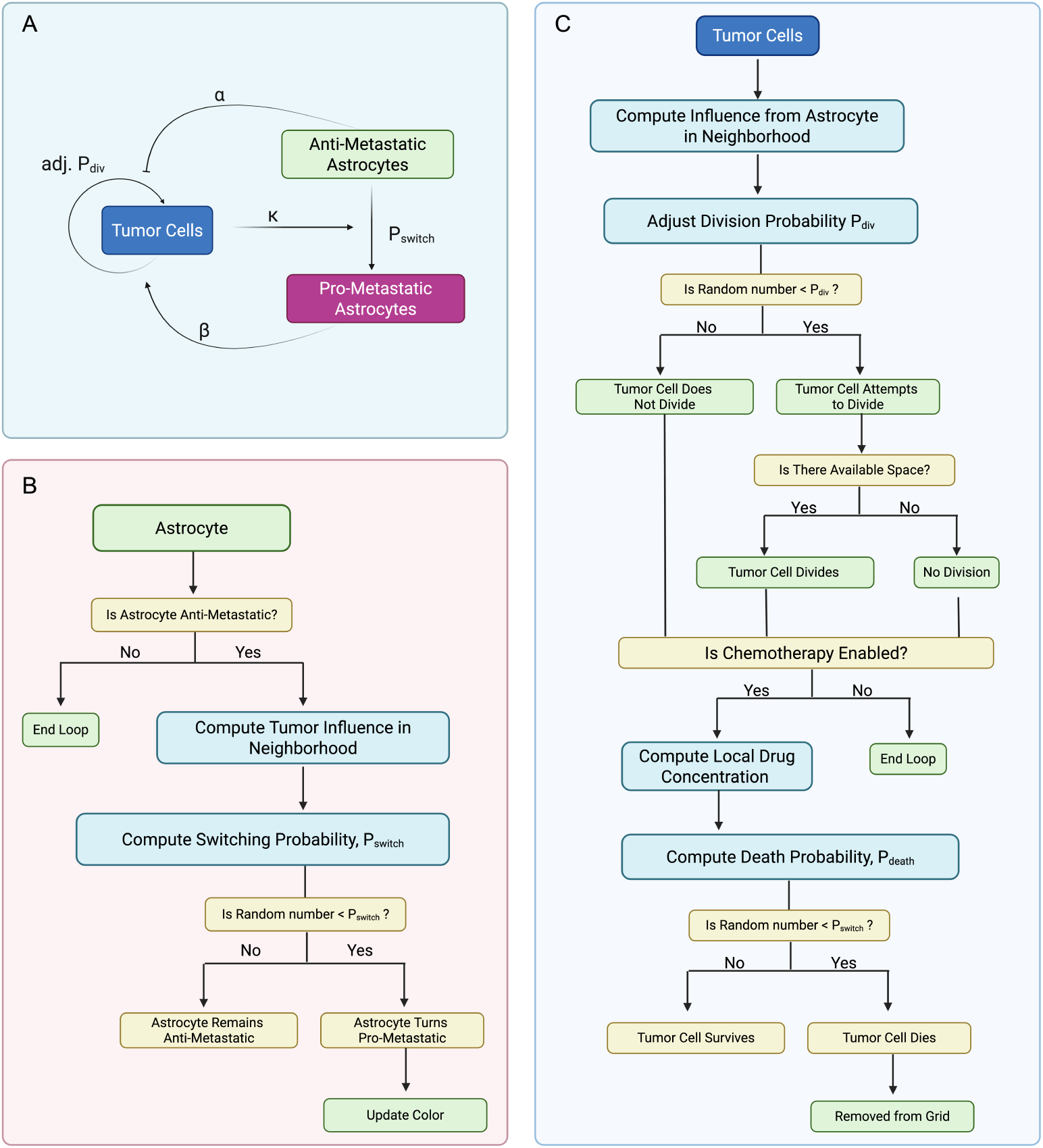
Agent-Based Model. (A) Schematic of cell–cell interactions. Tumor cells influence astrocyte state in a density-dependent manner. Anti-metastatic astrocytes reduce the rate of tumor cell proliferation (*α*), while pro-metastatic astrocytes enhance the rate of tumor cell proliferation (*β*). Astrocytes switch from anti-metastatic to pro-metastatic with probability *P*_switch_ based on the number of tumor cells present in the local neighborhood, per the astrocyte update loop. (B) Astrocyte update loop. Anti-metastatic astrocytes stochastically switch to a pro-metastatic state based on a probability influenced by the local tumor cell density. (C) Tumor cell update loop. Tumor cells adjust division probability based on local astrocyte influence, attempt division if a neighboring space is available, and undergo chemotherapy-induced death with probability *P*_death_ based on local drug concentration. *Created with BioRender*.*com*

**Fig. 2.**
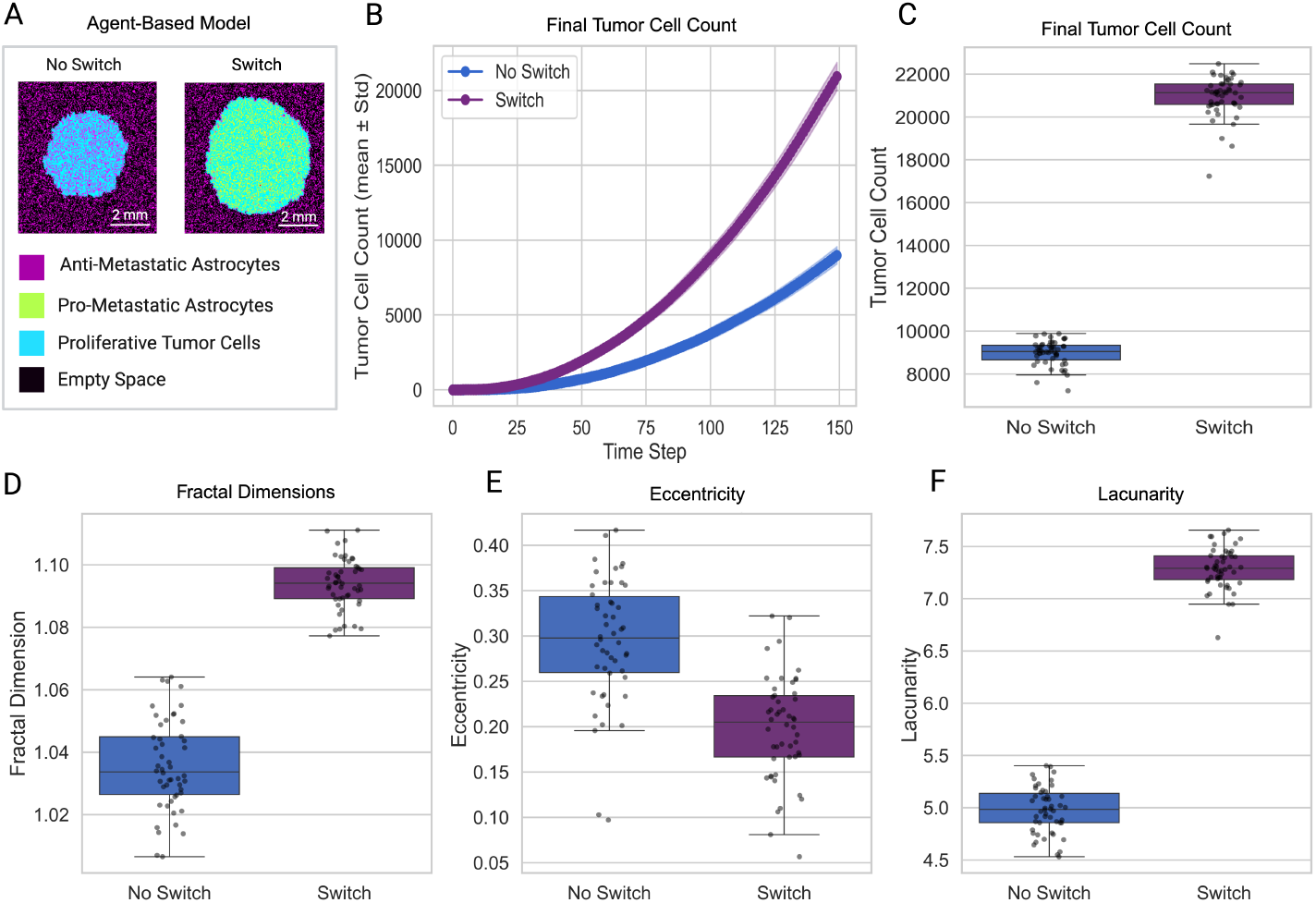
Astrocyte reprogramming enhances tumor growth and alters spatial morphology. (A) Representative agent-based model snapshots at the final time step show increased tumor size in the reprogrammable condition. (B) Tumor growth curves over time indicate that tumors in the reprogramming condition expand significantly faster than the non-reprogrammable condition. (C) Final tumor cell count distributions confirm a substantial increase in tumor burden with reprogramming. (D) Box plots of fractal dimension, eccentricity, and lacunarity at the final time step reveal that tumors with astrocyte reprogramming exhibit higher fractal dimension and increased lacunarity.

We quantified changes in spatial morphology using fractal dimension, lacunarity, and eccentricity (Fig. 2D-F). Tumors from simulations with astrocyte reprogramming showed slightly greater fractal dimension (Δ*F* ≈ +0.058, Mann–Whitney *U* = 0.0, *p* = 7.07 *×* 10^*−*18^), lower eccentricity (Δ*E* ≈ *−*0.096, Mann–Whitney *U* = 2173.0, *p* = 2.02 *×* 10^*−*10^), and higher lacunarity (ΔΛ ≈ +2.309, Mann–Whitney *U* = 0.0, *p* = 7.07*×*10^*−*18^). These changes reflect a shift towards increased boundary complexity and more rounded overall shape (reduced eccentricity) in the reprogrammed tumors. Tumors in the simulations with astrocyte reprogramming exhibited persistently higher lacunarity throughout the time course of the simulation, with separation from the group of simulations without astrocyte reprogramming emerging early and maintained throughout (Fig. S4). In contrast, eccentricity declined over time in both groups (Fig. S4).

To assess whether these morphological differences were driven by tumor size, we regressed out tumor cell count at each time step and computed residualized metrics. Although residual trajectories were more closely aligned across groups, statistically significant differences between tumors in the two groups remained at the final time step for both fractal dimension and lacunarity (Fig. S4). Specifically, residualized fractal dimension values were significantly elevated, while residualized lacunarity values were significantly reduced in the reprogrammable condition compared to the non-reprogrammable group (Mann–Whitney *U* = 184.0, *p* = 2.05 *×* 10^*−*13^; *U* = 1675.0, *p* = 3.43*×*10^*−*3^, respectively). Residualized eccentricity did not differ significantly between groups (*U* = 1135.0, *p* = 0.43). These results indicate that astrocyte reprogramming influences tumor spatial complexity and heterogeneity beyond effects attributable to overall tumor size. Notably, these differences were not driven by progressively diverging temporal trajectories but instead emerged as consistent, cumulative effects visible at the simulation endpoint.

### Astrocyte Suppression and Conversion Threshold Dominate Tumor Dynamics in Global Sensitivity Analysis

To assess whether differences in tumor size and spatial morphology persisted across parameter regimes, we performed a global sensitivity analysis using Sobol sequence–based sampling Renardy et al. (2021). Six model parameters—four continuous and two discrete—were varied over their expected biological ranges (Table 1). We observed that tumor burden was most sensitive to *θ* and *α*, both of which showed strong negative correlations, indicating that more stringent switching *θ* and anti-metastatic astrocyte influence *α* strongly suppress tumor growth. Conversely, *β* (effectProMet) exhibited a moderate positive correlation with tumor size, consistent with its growth-promoting role.

**Table 1.** Base parameters.

| Symbol | Parameter Name | Range | Biological/Model Meaning |
| --- | --- | --- | --- |
| $\alpha$ | effectAntiMet | 0–1 | Strength of tumor suppression by anti-metastatic astrocytes |
| $\beta$ | effectProMet | 0–1 | Strength of tumor promotion by pro-metastatic astrocytes |
| $\theta$ | conversionThreshold | 0–1 | Tumor influence level required for astrocyte state transition |
| $\kappa$ | effectPerTumorCell | 0–1 | Influence exerted per tumor cell on nearby astrocytes |
| $S_A$ | switchSensitivity | 8, 16, 32 | Steepness of the astrocyte state-switching probability curve |
| $S_T$ | divisionSensitivity | 4, 8, 16 | Sensitivity of tumor proliferation probability to astrocyte influence |

Fractal dimension and lacunarity were similarly sensitive to *θ* and *α*, with negative correlations indicating that suppressive astrocyte environments reduce boundary complexity (Fig. 3). Interestingly, *β* modestly increased lacunarity while having minimal effect on fractal dimension, suggesting that the tumor structure becomes more uneven locally with an increase in prometastatic effect, but without a corresponding increase in global boundary complexity. Eccentricity was most sensitive to *θ* and *α* in the positive direction, suggesting that these inhibitory mechanisms might promote elongated tumor morphologies. Other parameters, such as *S*_*A*_ (switchSensitivity) and *S*_*T*_ (divisionSensitivity), exhibited consistent but weaker effects across metrics.

**Fig. 3.**
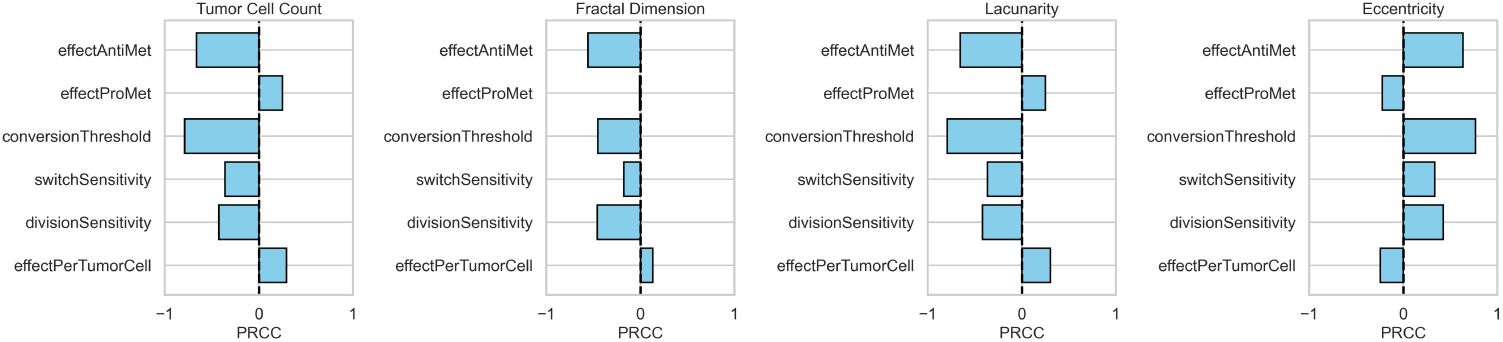
Partial rank correlation coefficients. Partial rank correlation coefficients (PRCC) between each model parameter and key output. Bars to the right (positive) indicate that higher parameter values increase the respective outcome, while bars to the left (negative) indicate an inverse relationship. Magnitude of each bar reflects the strength of the parameter’s influence.

To further contextualize these results, we examined the trends as each parameter was varied and key model outputs (tumor cell count, fractal dimension, lacunarity, and eccentricity). These analyses revealed notable variability within the outputs of certain parameters. For instance, tumor cell count variance increased with division sensitivity, rising from 3.84 *×* 10^7^ at *S*_*T*_ = 4.0 to 1.47 *×* 10^8^ at *S*_*T*_ = 16.0 (Fig. S5). This monotonic increase in variance suggests that higher division sensitivity amplifies variability in tumor burden, potentially reflecting nonlinear effects or higher-order interactions not fully captured by PRCC alone Renardy et al. (2021).

### Tumor Progression is Modulated by Differences in Astrocyte Density

To examine the impact of astrocyte density on tumor progression, we introduced neutral agents into the model while maintaining a fixed total of 45,000 cells. Astrocytes and neutral agents together occupied 50% of the grid, and the proportion of astrocytes within this compartment was systematically varied from 0% to 50%.

Tumor cell count increased monotonically with astrocyte density (analysis of variance [ANOVA], *p* = 5.57*×*10^*−*49^), with all pairwise comparisons statistically significant (Tukey’s honestly significant difference [HSD] test), except between 30% and 40%. To ensure appropriate test selection, we assessed normality and homogeneity of variance across density groups. ANOVA was applied to tumor cell count, eccentricity, and fractal dimension. Kruskal–Wallis was used for lacunarity due to violations of ANOVA assumptions. Lacunarity varied significantly with astrocyte density (Kruskal–Wallis, *H* = 75.71, *p* = 6.62 *×* 10^*−*15^), with Dunn’s post hoc tests identifying significant differences emerging beyond 20% astrocyte density. Eccentricity decreased slightly with increasing astrocyte density (ANOVA, *p* = 0.0047), though post hoc tests revealed that only the comparisons between 0% and 30%, and 0% and 40% reached statistical significance. In contrast, fractal dimension did not significantly differ across conditions (ANOVA, *p* = 0.745). Together, these results suggest that astrocyte density primarily modulates tumor burden and boundary lacunarity, while its effects on eccentricity are more modest, and its influence on boundary geometric complexity (fractal dimension) is minimal.

To evaluate how astrocyte density modulates tumor behavior across diverse parameter regimes, we performed a separate global sensitivity analysis at each astrocyte density level. To evaluate how astrocyte density modulates tumor behavior across diverse parameter regimes, we performed a separate global sensitivity analysis at each astrocyte density level, as described in the Methods. To facilitate comparison across conditions, we ranked parameter sets by mean tumor cell count at the 10% astrocyte density level and stratified them into tertiles, corresponding to three outcome-based categories: “Inhibitory,” “Neutral,” and “Promoting,” corresponding to increasing levels of tumor cell count. The parameter sets corresponding to inhibitory and promoting regimes in 10% density were applied to 50% density condition. Consistent with our expectations, the Promoting regime exhibited significantly higher tumor cell counts and more pronounced morphological indicators (e.g., increased fractal dimension) compared to those parameter sets from the Inhibitory regime (Fig. 4C). Notably, the differences between regimes were amplified at 50% astrocyte density, suggesting that higher astrocyte density may have a dual role in suppressing tumor growth in certain parameter regimens, and the opposite in those parameter sets predisposed to aggressive behavior.

**Fig. 4.**
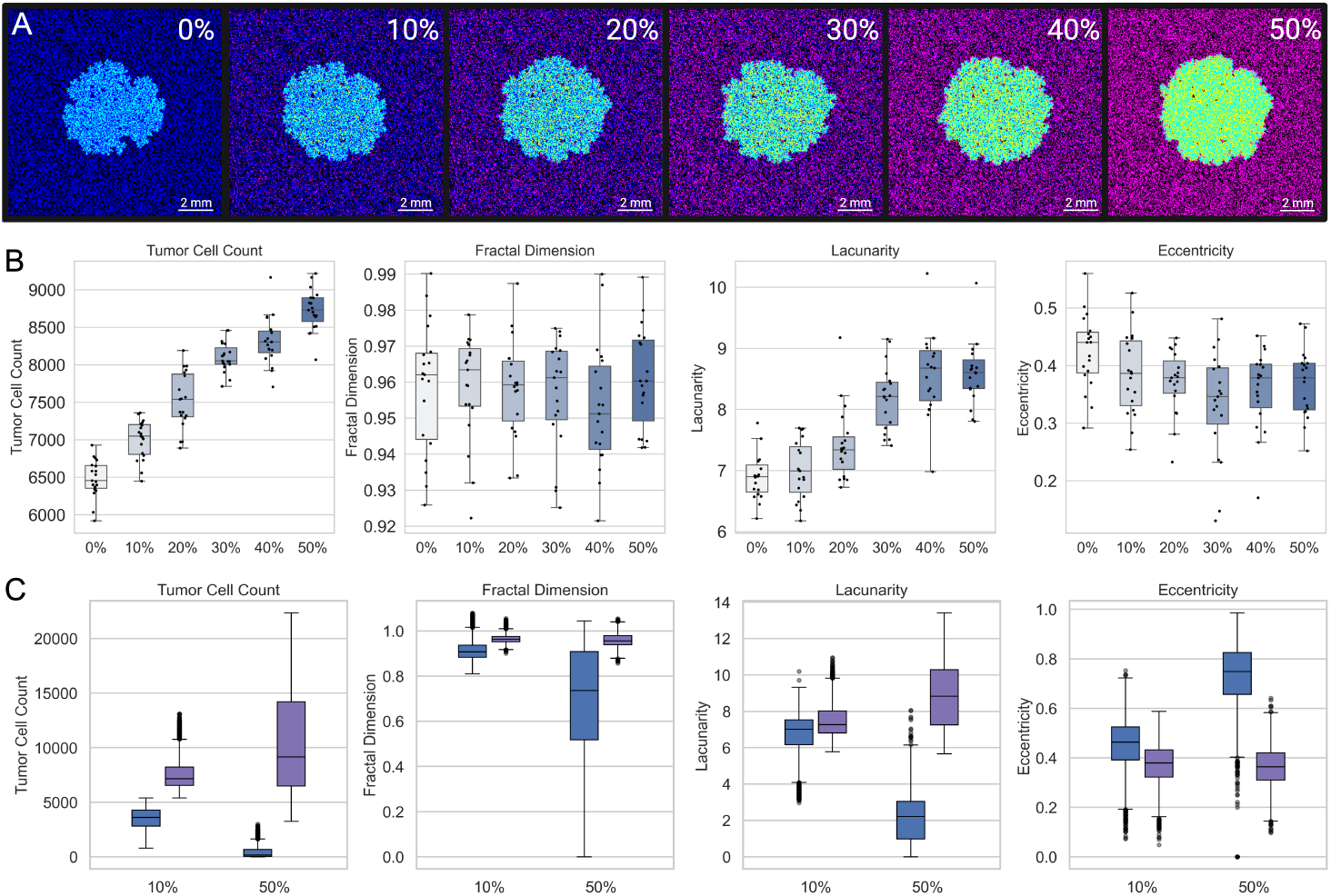
Effects of Astrocyte Density on Tumor Growth and Morphology. (A) Representative snapshots of the final tumor configuration at increasing astrocyte densities (0–50%). Astrocytes are initialized as anti-metastatic (purple) but turn pro-metastatic (yellow) with tumor cell (cyan) influence. Under higher astrocyte density, overall tumor burden decreases and spatial patterns become more heterogeneous. (B) Boxplots illustrating how tumor cell count, fractal dimension, lacunarity, and eccentricity vary with astrocyte density at the final time step. Higher astrocyte densities lead to reduced tumor sizes and alterations in tumor boundary morphology. (C) Boxplots comparing inhibitory and promoting regimes at two astrocyte densities (e.g., 10% and 50%).

### Astrocyte Spatial Distributions Drive Changes in Tumor Progression

We next evaluated how astrocyte spatial distribution affected tumor growth characteristics. We simulated tumor progression under six distinct spatial arrangements of astrocytes—uniform, random, clustered, radial, inverse radial, and gradient—while maintaining astrocyte density constant at 30% (Fig. 5A). Despite identical baseline parameters, the final tumor outcomes varied substantially across spatial configurations (Fig. S6). Because several groups violated normality or homogeneity of variance assumptions, we used the Kruskal–Wallis test followed by Dunn’s post hoc comparisons. Tumor cell count differed significantly across configurations (*p* = 1.19 *×* 10^*−*15^), with the inverse radial condition yielding the highest burden (23834 ± 579) and radial the lowest (9202 ± 1871). Fractal dimension also varied significantly across spatial patterns (*p* = 2.12 *×* 10^*−*14^), with uniform (1.116 ± 0.007) and inverse radial (1.111 ± 0.007) distributions associated with the highest values, while radial was lowest at 1.023 ± 0.030. Lacunarity differed across conditions as well (*p* = 1.36 *×* 10^*−*14^), decreasing from a high of 9.120 ± 0.407 in the clustered condition to 6.603 ± 0.477 in the radial condition, with the remaining distributions showing intermediate values. Notably, eccentricity varied considerably (*p* = 4.33 *×* 10^*−*11^): radial and gradient tumors exhibited the highest mean eccentricity (0.365±0.093 and 0.413±0.032, respectively), whereas uniform (0.172 ± 0.045) and inverse radial (0.178 ± 0.040) produced more isotropic tumor shapes. Together, these results demonstrate that spatial distribution of astrocytes – independent of overall density – can substantially influence tumor growth and morphology.

**Fig. 5.**
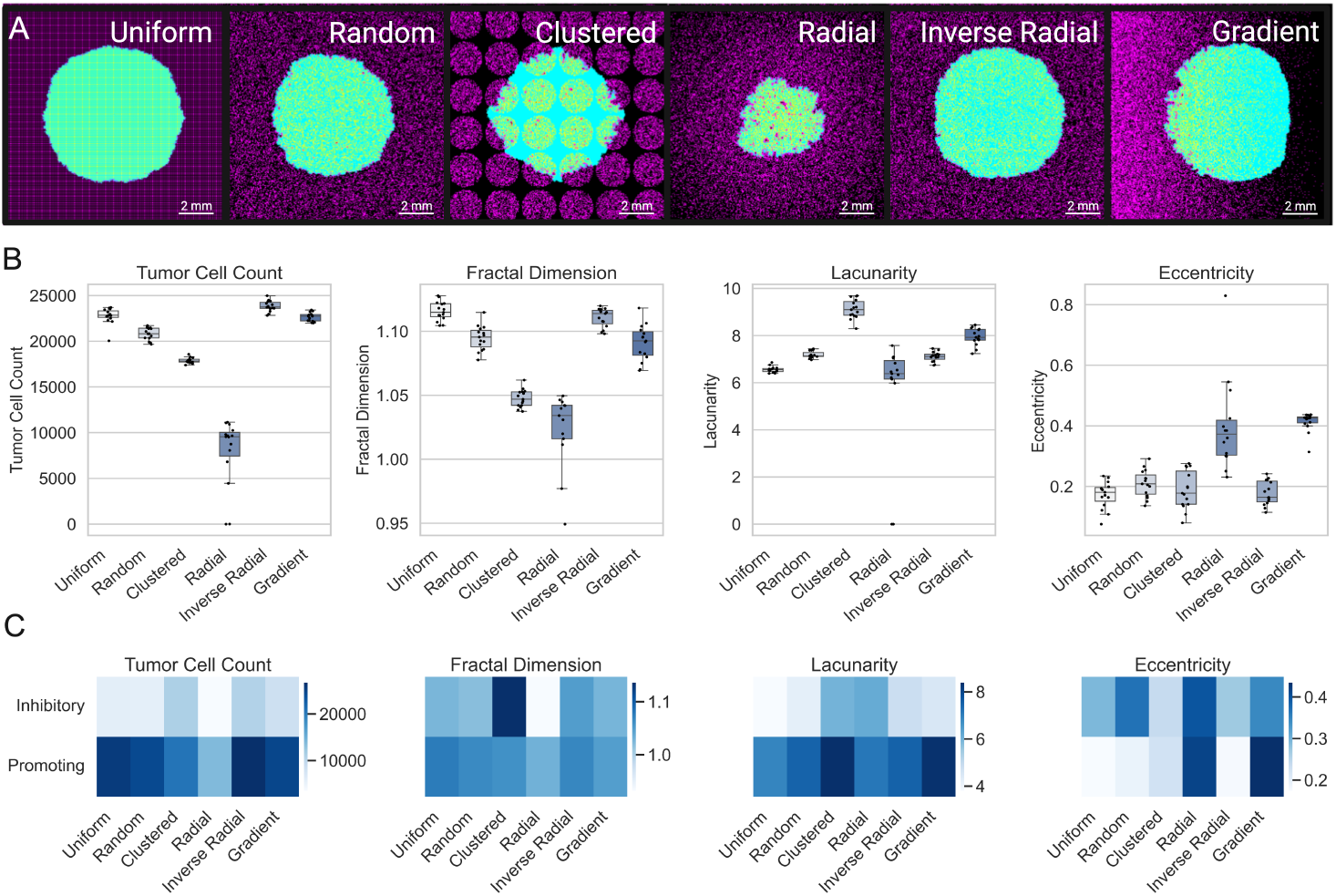
Effects of Astrocyte Spatial Distributions on Tumor Growth and Morphology. (A) Representative final snapshots from the agent-based model under six distinct astrocyte spatial arrangements (uniform, random, clustered, radial, inverse radial, gradient), with astrocytes shown in purple and tumor cells in cyan. All simulations use the same baseline parameters at 30% astrocyte density. (B) Boxplots of key outputs for each spatial distribution. (C) Heatmaps showing the mean final values of each metric across the six spatial distributions (columns) for parameter sets classified as either Inhibitory or Promoting (rows). Regime labels were derived by clustering on the uniform (grid 0) condition, and the same classification was applied to all distributions.

To evaluate how astrocyte spatial distributions modulated outcomes across different parameter regimes, we performed a global sensitivity analysis across all six spatial configurations. Mirroring our density-based analysis, we assigned parameter sets to Inhibitory, Neutral, and Promoting regimes based on final tumor cell count under the uniform distribution. The Inhibitory and Promoting regimes were then used to assess how tumor behavior varied across six distinct spatial configurations of astrocyte placement under different parametric conditions (Fig. 5C). Tumor burden and spatial morphology metrics (fractal dimension, lacunarity, and eccentricity) were summarized within each regime-configuration pair and visualized as heatmaps (Fig. 5C). Even under identical densities and parameter values, spatial configuration alone could tip the balance between tumor suppression and aggressive expansion (Fig. 5C). These findings support the hypothesis that tissue-level astrocyte heterogeneity may shape brain metastatic niche formation and progression through spatially constrained reprogramming dynamics.

### Astrocyte Reprogramming Confers Resistance to Chemotherapy *in silico*

In addition to providing proliferative advantages to tumor cells, astrocytes in our model confer localized chemotherapy resistance, consistent with observations of astrocyte–mediated chemoprotection in astrocyte-tumor co-cultures conducted *in vitro* Qu et al. (2023). This effect was implemented by reducing the effective drug concentration sensed by tumor cells based on the number of proximate pro-metastatic astrocytes, modulated by a gap junction modulation factor (*G*_*f*_) (Eq. 8). Although astrocytes do not absorb drug in our simulations, this modeled reduction in effective exposure reflects experimental findings that astrocyte proximity enhances tumor cell survival through gap junction signaling and paracrine support Qu et al. (2023). Higher *G*_*f*_ values lead to greater reductions in cytotoxic exposure, enabling tumor cells within astrocyte-rich regions to exhibit increased survival under periodic chemotherapy cycles.

Figure S7 illustrates these effects for three *G*_*f*_ values (0.0, 0.5, and 1.0), highlighting the spatial distribution of chemotherapy protection. At the lowest modulation (*G*_*f*_ = 0.0), overall tumor burden remains moderate, with only a few clusters showing heightened resistance. In contrast, when *G*_*f*_ = 1.0, direct tumor–astrocyte contact leads to near-complete drug blockade, forming extensive “hot spots” of high resistance (shown in purple) throughout the tumor core and resulting in a markedly larger final tumor mass.

We simulated tumor growth under periodic drug administration, with and without astrocyte reprogramming (N = 50) (Fig. 6A). In all simulations, drug treatment was initiated once the tumor cell population surpassed 3,000 cells, administered in 21-day cycles. Although each model initially underwent partial tumor regression or stabilization, the reprogrammable model exhibited a rapid rebound post-therapy, ultimately surpassing its baseline tumor size. By comparison, tumors without astrocyte reprogramming demonstrated prolonged periods of stagnation or decline with each treatment cycle (Fig. 6B). After two 21-day chemotherapy cycles, tumors in the strictly anti-metastatic simulations underwent a mean reduction to 1988 ± 822 cells, corresponding to a 34% decrease in tumor burden. In contrast, when astrocytes were allowed to switch states, tumors continued to expand and reached a mean size of 16,222 ± 652 cells. Pro-metastatic astrocytes thus create localized zones of reduced drug efficacy, enabling persistent tumor growth that resists multiple rounds of chemotherapy. These findings support the concept of microenvironment-mediated resistance driven specifically by reprogrammed astrocytes via gap junction crosstalk Chen et al. (2016).

**Fig. 6.**
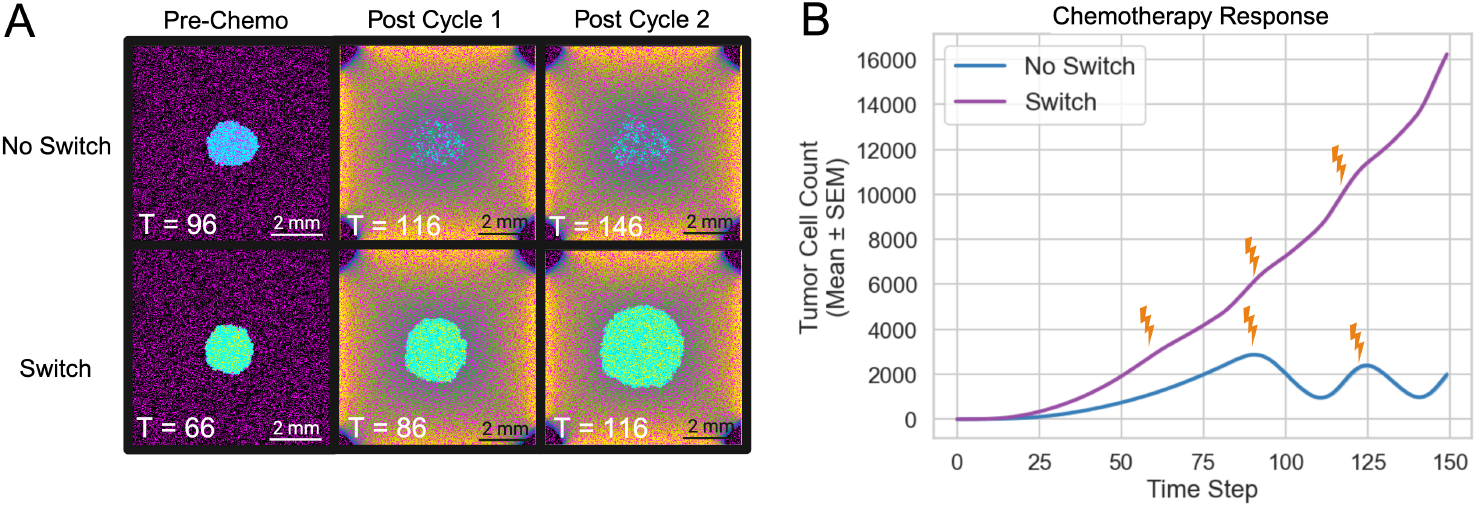
Chemotherapy Response. (A) Spatial snapshots at three key time points (Threshold, Cycle 1, and Cycle 2) for two model conditions: Anti-Metastatic Only (top row) and Reprogrammable (bottom row). The specific simulation times (T) are indicated on each snapshot. (B) Mean tumor cell count (± SEM) over time, averaged across 50 replicates, for each condition.

## Methods

### Model Overview

We developed a two-dimensional hybrid agent-based model (ABM) using the Hybrid Automata Library (HAL) framework to simulate tumor–astrocyte interactions within an avascular brain microenvironment Bravo et al. (2020). The model is implemented on a 300 *×* 300 lattice grid, where each grid site represents a 25 *×* 25 *µ*m region. Time is discretized into 17-hour time steps, chosen such that the probability of division per time step is approximately 0.5, assuming an average tumor cell doubling time of 34 hours Samson et al. (2021); Sweeney et al. (1998). Simulations are run for 150 time steps, corresponding to approximately 3.5 months of biological time. Each grid site can accommodate a single agent, either a tumor cell, an astrocyte, or a neutral agent. The model is initialized with a single tumor cell positioned at the center of the grid, with astrocytes distributed across the domain at varying densities and spatial configurations. All astrocytes are initialized in an anti-metastatic state but can transition to a pro-metastatic state in response to tumor-derived signals (Fig. 1A,B). This transition is modeled as an irreversible phenotypic switch, with the switching probability defined as a function of the local tumor cell density (Eq. 5, Fig. 7A,B). Once reprogrammed, pro-metastatic astrocytes enhance tumor cell proliferation by increasing the division probability of nearby tumor cells, while anti-metastatic astrocytes exert a suppressive influence (Eq. 10, Fig. 7A,C). All simulations are stochastic, with agent behaviors updated probabilistically at each discrete time step using a random sequential update scheme.

**Fig. 7.**
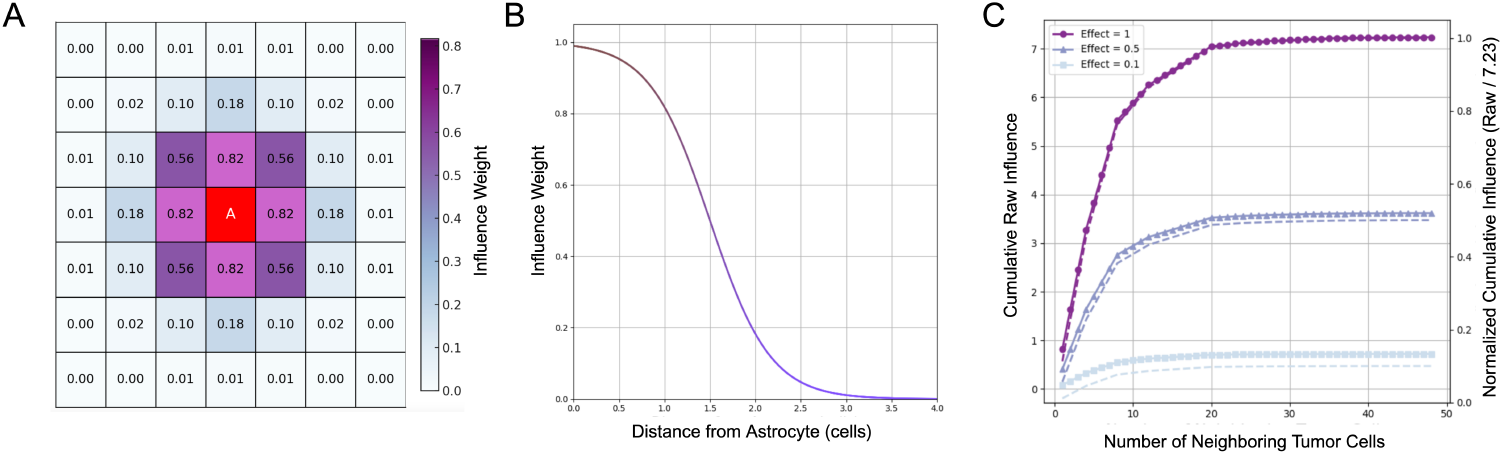
Astrocyte Reprogramming. (A) Schematic of the tumor cell influence on a central astrocyte based on Euclidean distance within a 7*×* 7 neighborhood. Each neighboring tumor cell contributes an influence value (*κ* = 1.0) calculated using a sigmoidal decay function with parameters *S* = 3 and *d* = 1.5. (B) Sigmoidal decay function describing how influence weight decreases with increasing distance from the astrocyte. The inflection point *d* = 1.5 corresponds to the distance at which the influence is reduced by half. (C) Cumulative tumor cell influence as a function of neighborhood size for three different per-cell effect magnitudes (*κ* = 1.0, 0.5, and 0.1). Solid lines show raw cumulative influence, while dashed lines represent normalized influence (raw / 7.23). The raw and normalized curves illustrate how local tumor density and per-cell effect magnitude modulate the astrocyte’s total exposure.

The model framework was also extended to account for the effects of treatment. Chemotherapy is modeled as a diffusible factor, governed by a reaction-diffusion partial differential equation:

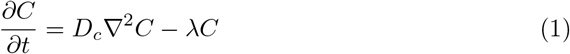

where, *D*_*c*_ is the diffusion coefficient and *λ* is the decay rate. Drug delivery was simulated by applying Dirichlet boundary conditions (*C* = *C*_*b*_) during dosing periods, representing peripheral drug influx. Between dosing periods, zero-flux (Neumann) boundary conditions 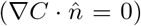 were applied to model restricted exchange at the brain boundary. Chemotherapy treatment is initiated when the tumor reaches a threshold of 3,000 cells, corresponding to a lesion approximately 1-1.5 mm in diameter in 2D space. This threshold reflects a micrometastatic tumor that is below clinical detection limits Yin et al. (2022) but large enough to evaluate treatment effect after initial tumor establishment.

Capecitabine, a fluoropyrimidine with modest brain penetration and with clinical use in brain metastases from triple-negative and HER2-positive breast cancer, was modeled as a representative chemotherapeutic agent Carvalho Gouveia et al. (2023); Morikawa et al. (2014). Treatment follows a 21-day cycle, with drug applied at each timestep for 14 consecutive steps, followed by a 7-step rest period Wagstaff et al. (2003). This regimen reflects standard clinical dosing (14 days on, 7 days off) and allows evaluation of tumor response under periodic therapeutic pressure and microenvironmental protection.

Tumor cell death is modeled as a function of local drug concentration (Eq. 13). Primary model outputs include final tumor cell count, fractal dimension, lacunarity, and eccentricity, which serve as quantitative metrics of tumor burden, spatial morphology, and therapeutic response. A full list of fixed model parameters and justifications is provided in Supplementary Table S1.

### Astrocyte Switch Modeling

We suppose that astrocyte reprogramming in BCBM is driven by sustained exposure to tumor-derived paracrine signals that promote a pro-metastatic phenotype Gong et al. (2019). To model this irreversible transition, we implemented a cumulative influence framework in which each astrocyte integrates input from tumor cells within a 7 *×* 7 square neighborhood (i.e., a 3-cell radius in grid units). Signal strength decays with distance according to a sigmoidal function and at each time step, total tumor influence is computed by summing the distance-weighted contributions from all nearby tumor cells (Fig. 7A). The effective influence of a tumor cell at a distance *d*_*i*_ from an astrocyte is given by:

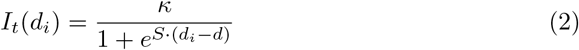

where, *κ* (effectPerTumorCell) denotes the magnitude of tumor influence on an astrocyte, *S* = 3 controls the steepness of decay over distance, and *d* = 1.5 denotes the half-maximal distance of suppression (Fig. 7B). The cumulative influence from all neighboring tumor cells surrounding an astrocyte is given by:

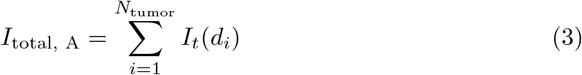

where *N*_tumor_ denotes the number of tumor cells within an astrocyte’s 7 *×* 7 neighborhood. To ensure compatibility across conditions, we normalize *I*_total, A_ by the theoretical maximum cumulative influence an astrocyte could experience,

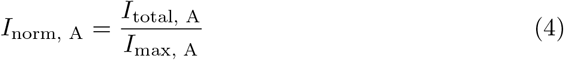

where *I*_max, A_ is the maximum possible sum of tumor cell influences in the local neighborhood (Fig. 7C). This normalization constrains *I*_norm_ ∈ [0, 1], corresponding to the absence and maximum presence of tumor influence, respectively. The probability of astrocyte state transition is then determined by a sigmoidal function:

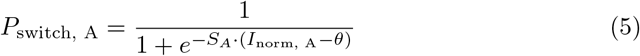

where *S*_*A*_ (switchSensitivity) controls the steepness of the transition curve, and *θ* (conversionThreshold) represents the normalized influence level at which the switching probability equals 50% (Fig. S1). At each time step, a random number is drawn from a uniform distribution. If this number is less than *P*_switch, A_, the astrocyte irreversibly switches to a pro-metastatic state.

### Tumor Cell Division

Tumor cell division is governed by a probability density function calibrated to yield a 50% division probability per time step in the absence of astrocyte influence, consistent with the observed average doubling time (∼34 hours) of MDA-MB-231 breast cancer cells Samson et al. (2021); Sweeney et al. (1998). In this model, anti-metastatic astrocytes reduce the probability of tumor proliferation, while pro-metastatic astrocytes increase the probability of tumor cell proliferation. Although astrocyte-mediated suppression is primarily pro-apoptotic *in vivo*, we model its net effect as reduced proliferation to simplify implementation while preserving its impact on tumor growth dynamics Valiente et al. (2014). We model the influence of an anti-metastatic astrocyte *j* at a distance *d*_*j*_ from a tumor cell using a sigmoidal decay function:

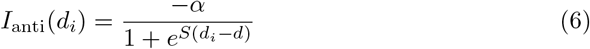

where *α* (effectAntiMet) represents the magnitude of tumor division suppression exerted by anti-metastatic astrocytes, *S* = 3 controls the steepness of decay over distance, and *d* = 1.5 denotes the half-maximal distance of suppression. The influence of pro-metastatic astrocytes on tumor cell proliferation is similarly defined:

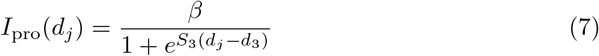

where *β* (effectProMet) represents the magnitude of tumor division promotion exerted by each pro-metastatic astrocyte. *S* and *d* retain the same values as in Equation (1). The total influence of all astrocytes within a 7 *×* 7 square neighborhood of a tumor cell is given by:

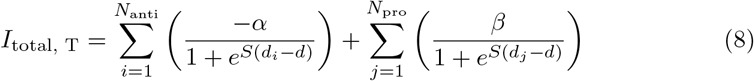

where *N*_anti_ and *N*_pro_ represent the number of anti-and pro-metastatic astrocytes, respectively. To constrain this value to a standardized range, the influence is normalized:

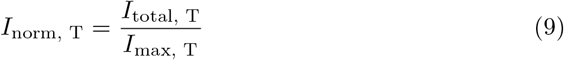

where *I*_max, T_ is the maximum theoretical cumulative astrocyte influence, ensuring *I*_norm, T_ ∈ [*−*1, 1]. The final division probability is given by a sigmoidal function:

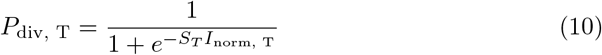

where *S*_*T*_ (divisionSensitivity) controls the sensitivity of tumor cell division to astrocyte influence. When *I*_norm, T_ = 0, *P*_div, T_ = 0.5, preserving baseline proliferation in the absence of astrocytes (Fig. S1). At each time step, a tumor cell attempts division if a random draw from a uniform distribution is less than *P*_div, T_ and if an empty space is available.

### Chemotherapy Implementation

Chemotherapy is introduced once the tumor population reaches a predefined threshold of 3,000 cells. Treatment is administered in 21-day cycles, with drug exposure applied at each timestep for 14 consecutive days, followed by a 7-day rest period Wagstaff et al. (2003). The drug is applied at the grid edges and diffuses inward following an Alternating Direction Implicit (ADI) diffusion scheme. Absorbing boundary conditions ensure that any drug molecules diffusing beyond the simulation domain do not reenter. The local chemotherapy concentration, denoted as *C*(*x, y, t*) at grid position (*x, y*) and time *t*, is updated each timestep according to:

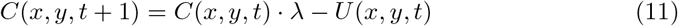

where, *λ* represents the chemotherapy decay rate and *U* (*x, y, t*) denotes the amount of drug absorbed by tumor cells, modeled as a fixed fraction of the drug available at a cell’s location.

To capture the chemoprotective effect of astrocyte gap junction communication, we modulate the effective local drug concentration around tumor cells based on the number of neighboring pro-metastatic astrocytes in their direct Moore neighborhood. If a tumor cell is surrounded by *N*_pro_ pro-metastatic astrocytes (out of a maximum of 8), the effective concentration is given by:

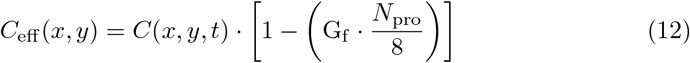

where *G*_*f*_ quantifies the maximal fractional reduction in drug efficacy due to gap junction signaling and *C*_eff_ (*x, y*) is the effective drug concentration detected by a tumor cell. Tumor cell death due to chemotherapy is modeled using a sigmoid function of the effective local drug concentration. For each tumor cell located at (*x, y*), the death probability *P*_death_ per timestep is given by:

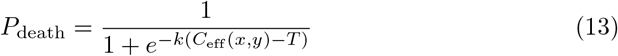

where *k* is the steepness parameter and *T* is the threshold concentration at which there is a 50% chance of death. At each timestep, an independent stochastic decision is made for each tumor cell: a random number is drawn from a uniform distribution, and if the number is less than *P*_death_, the tumor cell undergoes apoptosis and is removed from the grid. Chemotherapy does not affect astrocytes in our model, as their proliferation rate is significantly slower than that of tumor cells Colodner et al. (2005); Miyake et al. (1992). Given that chemotherapy primarily targets rapidly dividing cells, astrocytes are assumed to be resistant.

### Spatial Features

To quantify the morphological characteristics of tumor growth, we computed three spatial features: fractal dimension, lacunarity, and eccentricity. Fractal dimension and lacunarity are derived from the tumor boundary, defined as the outermost tumor cells at each time step. In contrast, eccentricity was calculated using the entire bulk tumor, which included all tumor cells within the growing mass. Tumor front cells, used for boundary-based calculations, were identified as tumor cells adjacent to at least one empty grid space within a 3 *×* 3 Moore neighborhood. This classification was validated through spatial visualization (Fig. S2), to ensure that only edge-proximal tumor cells were included in the boundary definition.

#### Fractal Dimension

Fractal dimension (*D*_*f*_) characterizes the complexity of the tumor boundary and was estimated using the box-counting method Curtin et al. (2021); Battalapalli et al. (2023); Iftekharuddin et al. (2003). At each time point, the tumor boundary was overlaid with a series of non-overlapping square grids of decreasing box sizes (*ϵ*). For each grid scale, the number of occupied boxes *N* (*ϵ*) was counted. The fractal dimension was computed as the slope of the log–log relationship:

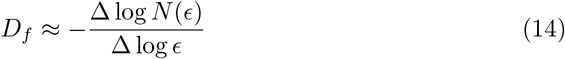

The slope was estimated using linear regression in log–log space. Higher *D*_*f*_ values indicate increased morphological complexity at the tumor boundary.

#### Lacunarity

Lacunarity (Λ) quantifies spatial heterogeneity by measuring the distribution of empty spaces within the tumor boundary Curtin et al. (2021); Plotnick et al. (1993). For each window size *ϵ* (i.e., a square region of side length *ϵ*), we partitioned the tumor grid into non-overlapping windows and, for each window, counted the number of grid sites containing tumor cells. This yielded a distribution of tumor counts across all windows, referred to as the mass distribution. Lacunarity was then defined as:

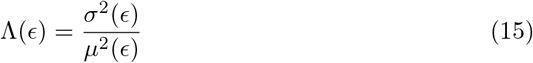

where *σ*^2^(*ϵ*) and *µ*(*ϵ*) are the variance and mean of the mass distribution, respectively. Higher lacunarity values indicate greater spatial irregularity at the tumor’s boundary.

#### Eccentricity

Eccentricity (*E*) quantifies the deviation of the tumor shape from circularity and is used to measure anisotropy of the tumor mass Mayer et al. (2021); Ismail et al. (2018). The tumor centroid was computed, and the principal axes were derived from the covariance matrix of the tumor cell coordinates. The corresponding eigenvalues were extracted to define the semi-axis lengths. Eccentricity was then calculated as:

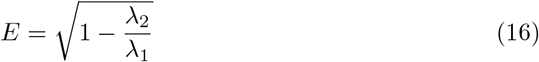

where *λ*_1_ and *λ*_2_ are the major eigenvalues of the covariance matrix, with *λ*_1_ ≥ *λ*_2_. A value of *E* ≈ 0 indicates a nearly circular tumor morphology, while values approaching 1 correspond to an elongated or irregular tumor shape.

## Astrocyte Heterogeneity

Astrocyte are initialized in the model with varying densities and spatial distributions to investigate their impact on tumor progression.

### Astrocyte Density

We define astrocyte density as the proportion of the simulation grid occupied by astrocytes at the beginning of the simulation, ranging from 0% to 50% of total lattice sites. Higher densities reflect astrocyte-rich brain regions, such as the cerebellum and hippocampus, while lower densities correspond to more sparsely populated regions like the cerebral cortex Man et al. (2024); Endo et al. (2022); Forrest et al. (2022).

To isolate the functional effects of astrocytes from overall cellular occupancy, the total number of non-tumor agents (astrocytes plus neutral agents) was fixed at 45,000 in all conditions, corresponding to 50% of the grid. In lower-density conditions, neutral agents were introduced to fill the remaining grid positions. These neutral agents do not interact with tumor cells, do not undergo state transitions, and are unaffected by any treatment or drug exposure modeled in the simulation. They serve solely to preserve constant total cell occupancy across conditions. This design ensures that any observed differences in tumor dynamics are attributable specifically to astrocyte function rather than to changes in cell crowding or diffusional constraints.

### Astrocyte Spatial Distributions

We modeled six different astrocyte spatial distributions to assess how initial placement patterns affect tumor growth and morphology: uniform, random, clustered, inverse-radial, radial, and gradient (Fig. 8). These patterns were chosen to represent a spectrum of biological realism and theoretical controls. The random, inverse-radial, and gradient distributions reflect biologically plausible arrangements. For example, the inverse-radial distribution mimics histological observations in which astrocytes are enriched at the tumor periphery, while the gradient distribution simulates the gray-white matter junction, where astrocyte density transitions gradually. The random distribution captures the dispersed organization of astrocytes in much of the brain parenchyma. In contrast, uniform, clustered, and radial patterns serve as theoretical constructs that help define the range of possible spatial effects. Astrocyte positions remain static throughout each simulation to isolate the impact of initial spatial structure on tumor growth and spatial morphology.

**Fig. 8.**
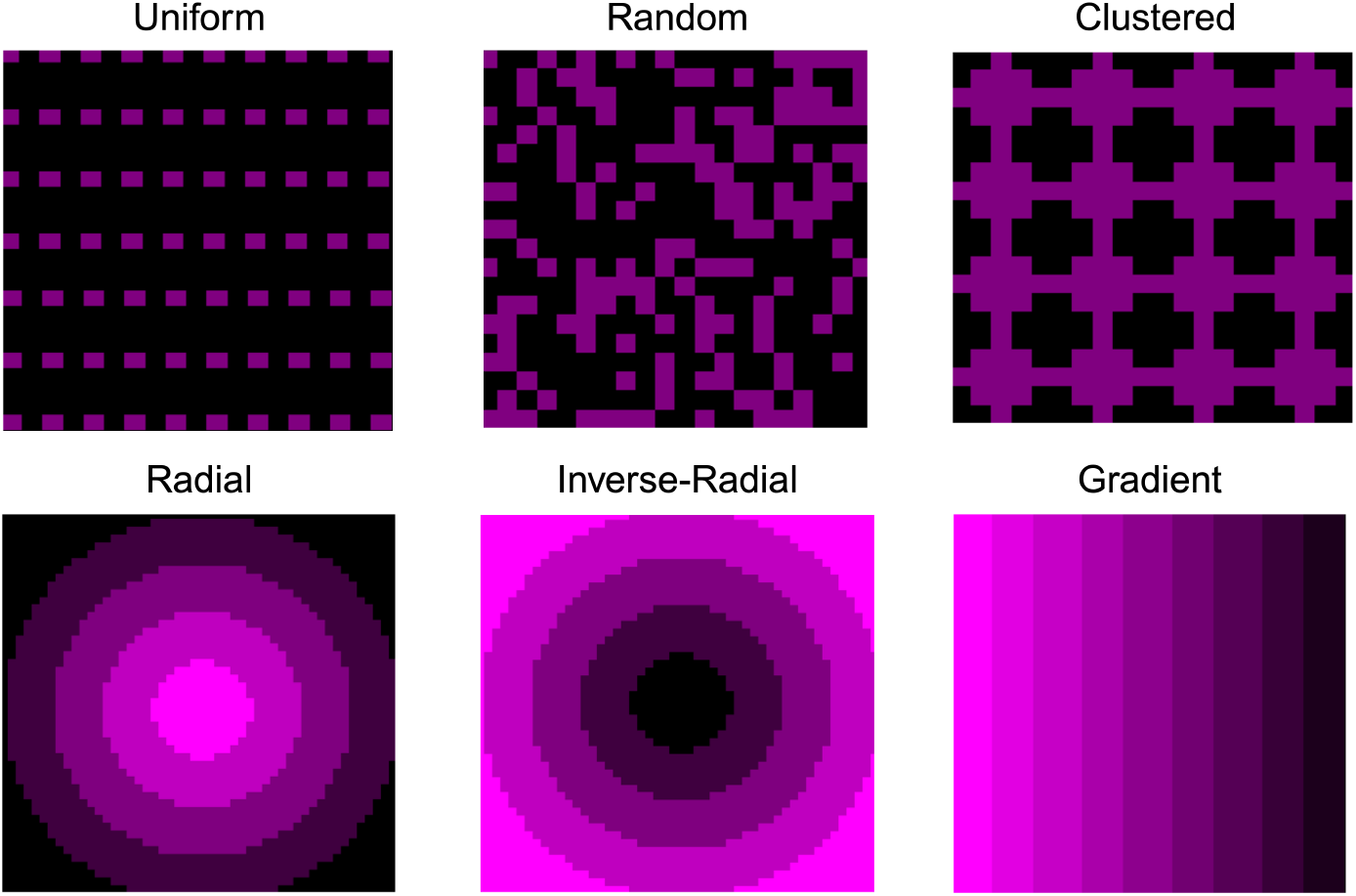
Spatial Distributions. (A) Uniform - Evenly spaced astrocytes (purple) with minimal spatial heterogeneity. (B) Random - Stochastically placed astrocytes, mimicking their broad distribution in brain parenchyma. (C) Clustered - Uniformly placed aggregated astrocytes forming localized clusters. (D) Radial - Astrocytes concentrate at the tumor core. (E) Inverse-Radial - Astrocyte density increases outward from the tumor. (F) Gradient - Astrocytes are arranged along a linear horizontal gradient.

### Sensitivity and Parameter Analysis

To evaluate the robustness of model predictions and identify critical parameters that govern tumor behavior, we performed a global sensitivity analysis. For the base-line model, we varied six core parameters governing astrocyte–tumor interactions (Table 1). Each parameter was sampled across a biologically plausible range using a Sobol sequence design (*n* = 1, 000), ensuring comprehensive coverage of the multi-dimensional parameter space (Fig. S3). We simulated 10 independent replicates per parameter set to account for stochastic variability in tumor growth. To quantify the influence of each parameter on model behavior, we computed partial rank correlation coefficients (PRCCs) between parameter values and key outputs.

We conducted two additional parameter sweeps with astrocyte density and spatial heterogeneity as independent variables, in addition to the six key variables. Each extended sweep included 1,000 parameter sets with 10 replicates per set, yielding 10,000 simulations per condition. The astrocyte density sweep tested five discrete density levels (0%, 12.5%, 25%, 37.5%, and 50%). The spatial distribution sweep included six predefined spatial patterns: uniform, random, clustered, inverse-radial, radial, and gradient.

## Discussion

Here, we present a novel, spatially explicit, two-dimensional agent-based model examining the role of astrocyte reprogramming in breast cancer brain metastases (BCBM). Our model simulates tumor–astrocyte interactions by allowing astrocytes to switch from an antito a pro-metastatic phenotype in response to local tumor cell density. By systematically varying astrocyte density, spatial distributions, and chemotherapy conditions, we characterized microenvironmental determinants of tumor progression and drug resistance. This work generates mechanistic hypotheses that can be tested in co-culture or *in vivo* models.

We found that astrocyte reprogramming substantially increases tumor burden and modifies tumor architecture. Tumors with reprogrammed astrocytes reached nearly 2.3 times the final size of those with astrocytes that were only simulated as anti-metastatic, and exhibited increased boundary fractal dimension and lacunarity, suggesting an irregular, spatially complex morphology, generally consistent with more invasive, heterogeneous growth. These morphological changes were not solely attributable to differences in tumor size. Our *in silico* findings reinforce experimental evidence that astrocytes modulate tumor burden Valiente et al. (2014); Kim et al. (2011) and further suggest that they can alter tumor spatial organization. While direct evidence in brain metastases is limited, co-culture models in glioma, breast and lung cancer indicate that astrocyte–tumor interactions can drive aggressive behavior Gagliano (2009); Zhang et al. (2015); Kim et al. (2011).

Our simulations also show that greater astrocyte densities in pro-metastatic regimes substantially amplify tumor burden and boundary irregularities, indicating that astrocyte-rich environments can act as niches permissive to metastatic expansion. Notably, spatial arrangement further modulates these outcomes: radial or uniform astrocyte distributions consistently yield larger, morphologically complex tumors, whereas inverse radial and clustered configurations sometimes constrain growth. These observations underscore the importance of both astrocyte density and local organization—beyond mere phenotypic state—in shaping the trajectory of brain metastatic colonization and invasive fronts.

From a clinical perspective, these results imply that anatomical context may be a key determinant of metastatic aggressiveness and influence treatment response. In certain brain compartments, astrocytes may be more densely concentrated or pre-disposed to adopt a pro-metastatic phenotype, and thus become “hot spots” for aggressive tumor expansion and early therapeutic failure. Moreover, given astrocytes’ role in promoting or maintaining metastatic dormancy, regional heterogeneity in astrocyte distribution may help explain why metastases preferentially colonize specific brain areas, while others remain refractory Quattrocchi et al. (2012). Although this hypothesis remains untested in brain metastases, it is consistent with broader evidence from glioma studies, where tumor location—and by extension, local microenvironment—might associate with tumor molecular subtype and clinical aggressiveness Altieri et al. (2017).

While astrocyte density can be estimated from reference atlases or regional transcriptomic data, spatial arrangement is typically unknown in human tissue and may vary stochastically. Our results suggest that such variation alone, even under fixed density and phenotypic conditions, can substantially alter tumor burden and morphology. Spatial organization should therefore be treated as a critical but often unobserved microenvironmental variable when linking cell–cell interactions to tumor architecture and behavior in patient data.

Our model also predicts that astrocyte reprogramming impairs the efficacy of cytotoxic chemotherapy. Tumors with reprogrammable astrocytes exhibited attenuated drug response and greater post-treatment regrowth, underscoring the limitations of conventional therapies in the context of a reactive microenvironment. These findings highlight astrocytes as a compelling therapeutic target. Unlike tumor cells, astrocytes are genetically stable and may be less prone to rapid resistance Quail and Joyce (2013). Several repurposed agents that inhibit astrocyte-mediated signaling—such as the endothelin receptor antagonist macitentan and gap junction inhibitor meclofenamate—are currently being evaluated in clinical trials, though they lack astrocyte specificity and may exert parallel effects on tumor or endothelial cells. Lee et al. (2016); Boire et al. (2017); Wasilewski et al. (2017). These efforts reflect both the promise and the current limitations of targeting astrocyte–tumor crosstalk. A more refined understanding of the transcriptional and genetic basis of astrocyte reprogramming may reveal more selective and durable intervention points.

While our model parameters were selected based on theoretical considerations and available experimental literature, the precise biological mappings are inherently approximate due to the lack of *in vivo* data. Despite this, the qualitative behaviors observed in our model—namely, spatially driven astrocyte reprogramming and its effects on tumor burden and morphology—emerge robustly across a wide range of biologically plausible parameter sets, supporting the generality of these dynamics.

Future extensions of this model could incorporate additional cellular components of the brain tumor microenvironment. For example, microglia and T cells are known to shape tumor–immune interactions and may also influence tumor progression through cytokine signaling or tissue remodeling Matias et al. (2018); Mirzaei and Yong (2022);

Feng et al. (2024). Integrating such cell types would allow for a more complete representation of brain metastatic ecology. Moreover, integration with spatially resolved transcriptomic data and histological imaging could provide parameter constraints and allow for *in silico* prediction of patient-specific progression trajectories Heindl et al. (2015).

In summary, our findings demonstrate that astrocyte reprogramming shapes both the kinetics and spatial morphology of breast cancer brain metastases, with direct implications for therapeutic response. We therefore show how, crucially, microenvironmental plasticity is a mediator of phenotypic heterogeneity and therefore therapeutic response in BCBM—contributing to variable treatment outcomes even in genetically similar tumors. Given the poor prognosis and limited efficacy of current therapies in this setting, strategies aimed at disrupting stromal reprogramming hold promise for improving patient outcomes in this challenging clinical context.

## Author contributions. Rupleen Kaur

Conceptualization; Formal analysis; Investigation; Methodology; Software; Visualization; Writing—original draft; Writing—review and editing. **Rowan Barker-Clarke**: Conceptualization; Supervision; Writing—review and editing. **Andrew Dhawan**: Conceptualization; Investigation; Supervision; Writing—review and editing.

## Data availability

All datasets and code used in this study will be made publicly available via GitHub upon publication.

