## Supplementary File for "Astrocyte reprogramming drives tumor progression and chemotherapy resistance in agent-based models of breast cancer brain metastases"

### Supplementary Figures

**Table S1 Fixed Model Parameters and Rationale**

| Parameter | Symbol | Value/Range | Rationale |
|---|---|---|---|
| **Spatial Scaling** | | | |
| Grid spacing | – | 25 $\mu$m | Represents a single cell diameter; establishes spatial resolution of the grid. |
| Grid size | – | 300 × 300 | 7.5 mm × 7.5 mm tumor field; balances resolution and computational cost. |
| Time step | – | 17 hours | Calibrated to MDA-MB-231 proliferation rate; one cell cycle spans two time steps. Samson et al. (2021); Sweeney et al. (1998) |
| **Drug Kinetics** | | | |
| Boundary drug conc. | $C_b$ | 0.095 | Sets outer boundary concentration for diffusion; scaled to match decay rate and maintain gradient. |
| Drug diffusion rate | $D_c$ | 0.979 $\text{mm}^2/\text{day}$ | Simulates realistic tissue-scale diffusion across 24 h. |
| Drug decay rate | $\lambda$ | 0.90 | Tuned to enable near-complete clearance between 21-day cycles, consistent with capecitabine kinetics. Reigner et al. (2001) |
| Drug absorption rate | $U$ | $-0.005$ | Small per-timestep uptake; preserves concentration gradient across tumor. |
| Drug modulation factor | $G_f$ | 0.3 | Models astrocyte-mediated resistance; attenuates without complete shielding. |
| **Sigmoid Response Properties** | | | |
| Sigmoid decay steepness | $S$ | 3 | Steepness of spatial decay for cell influence; selected to restrict interactions to local neighborhood. |
| Half-max decay distance | $d$ | 1.5 | Distance at which influence drops to 50%; sets neighborhood size for interaction functions. |
| **Drug Effect Parameters** | | | |
| Drug death threshold | $T$ | 0.5 | Drug concentration at which tumor cells have a 50% chance of death per timestep; selected to yield near-complete tumor response in the absence of astrocyte-mediated resistance. |
| Drug death steepness | $k$ | 5 | Controls steepness of death probability response; tuned to ensure a sharp transition in drug efficacy consistent with a cytotoxic agent. |

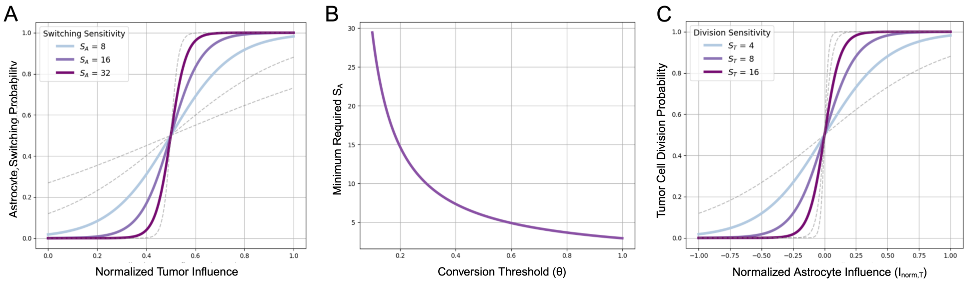


**Fig. S1 Sigmoid sensitivity functions for astrocyte switching and tumor proliferation.** (A) Sigmoid functions describing the probability of astrocyte state switching as a function of normalized tumor influence for varying values of astrocyte switching sensitivity ($S_A$). Higher $S_A$ yields a steeper switching curve, producing more abrupt transitions between anti- and pro-metastatic states. (B) Minimum $S_A$ required to ensure the switching probability $P(0)$ remains below 0.05, as a function of the conversion threshold $\theta$. (C) Sigmoid curves for tumor cell division probability as a function of normalized astrocyte influence ($I_{\mathrm{norm,T}}$) under varying tumor cell division sensitivity values ($S_T$). Increasing $S_T$ sharpens the response of proliferation probability to changes in local astrocyte state.

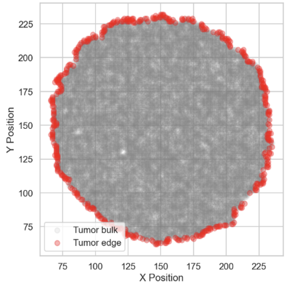


**Fig. S2 Tumor edge detection for spatial morphology analysis.** Visualization of simulated tumor configuration showing bulk tumor cells (gray) and edge cells (red). Tumor edge cells are identified as those with at least one adjacent empty site within their Moore neighborhood. This classification is used to define the tumor boundary for calculating spatial morphology metrics such as fractal dimension and lacunarity.

### Spatial Distribution Implementation Details

To evaluate how spatial heterogeneity in astrocyte positioning influences tumor progression, we implemented six distinct initial spatial distributions in the simulation grid.

- **Uniform:** Astrocytes were placed in an evenly spaced square grid to minimize spatial heterogeneity. The number of rows and columns was determined from the total astrocyte count to maintain near-equal spacing. Grid coordinates were computed using a regular mesh, and astrocytes were positioned at the center of each grid cell. The center position was excluded to avoid overlap with the tumor seed.
- **Random:** Astrocytes were placed stochastically across the grid using a uniform random distribution, subject to exclusion of the tumor center and already-occupied grid sites. Each unoccupied site had equal probability of being chosen.
- **Clumped:** The grid was first partitioned into a grid of rectangular blocks. Each block contained a single circular clump of radius $R$ (specified by `clumpRadius`) centred at the geometric centre of the block; if this coincided with the tumor seed, the centre was shifted by one lattice unit to avoid overlap. Within every clump, astrocytes were randomly placed over the grid. The target number of astrocytes, $N_{\text{astro}}$, was divided as evenly as possible among the clumps; any remainder was distributed by adding one extra astrocyte to the first few clumps. Candidate sites already occupied or lying outside the grid bounds were rejected.
- **Radial:** Astrocytes were distributed according to a weighted radial gradient centered on the tumor seed, with higher probabilities assigned to grid locations closer to the center. The steepness of the gradient was controlled by a user-defined exponent and scaling factor. The tumor site was excluded, and sampling was performed without replacement.
- **Inverse Radial:** Astrocytes were distributed based on their distance from the tumor seed, with higher placement probabilities assigned to peripheral grid locations. The

center of the grid was explicitly excluded to preserve the tumor site. The spatial bias toward the periphery was controlled by an exponent parameter, and astrocytes were selected without replacement from unoccupied sites according to these weights.

- **Gradient:** Astrocytes were placed according to a vertical gradient that favored the left edge of the grid. Each available site was assigned a weight based on its vertical position, with higher weights given to sites closer to the left edge. The steepness of the gradient was controlled by an adjustable exponent. The tumor seed site was excluded, and astrocytes were sampled without replacement based on the resulting spatial weights.

All spatial distributions were implemented in custom Java code. For weighted placement schemes, sampling was performed without replacement, with each unoccupied grid site assigned a probability proportional to its spatial weight. The tumor seed location was excluded from all configurations. To ensure comparability across conditions, the total number of astrocytes was held constant.

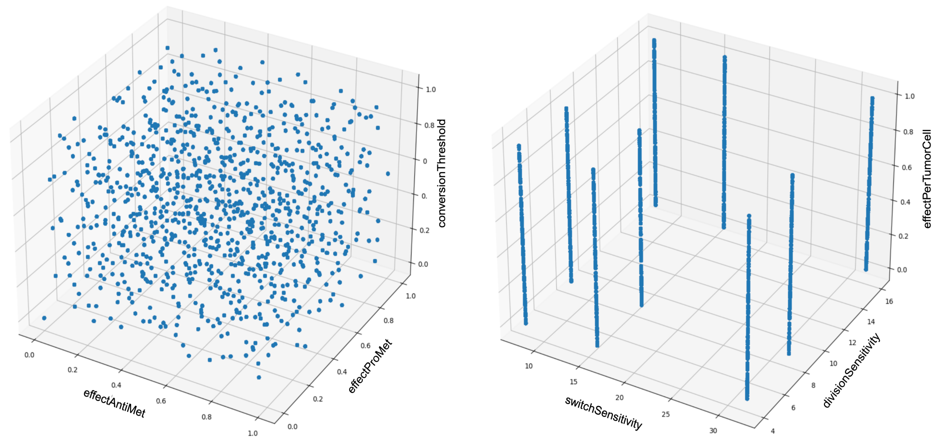


**Fig. S3 Sampling structure for continuous and discrete parameters in the sensitivity analysis.** (A) 3D scatter plot showing uniform sampling of three continuous parameters effectAntiMet ($\alpha$), effectPerTumorCell ($\kappa$), and conversionThreshold ($\theta$)—using Sobol sequence sampling to ensure space-filling coverage of the parameter space. (B) Discrete sampling of astrocyte switching sensitivity ($S_A$) and tumor division sensitivity ($S_T$), with values constrained to a predefined set (e.g., 8, 16, 32 for $S_A$ and 4, 8, 16 for $S_T$), visualized as vertical slices in the 3D parameter space.

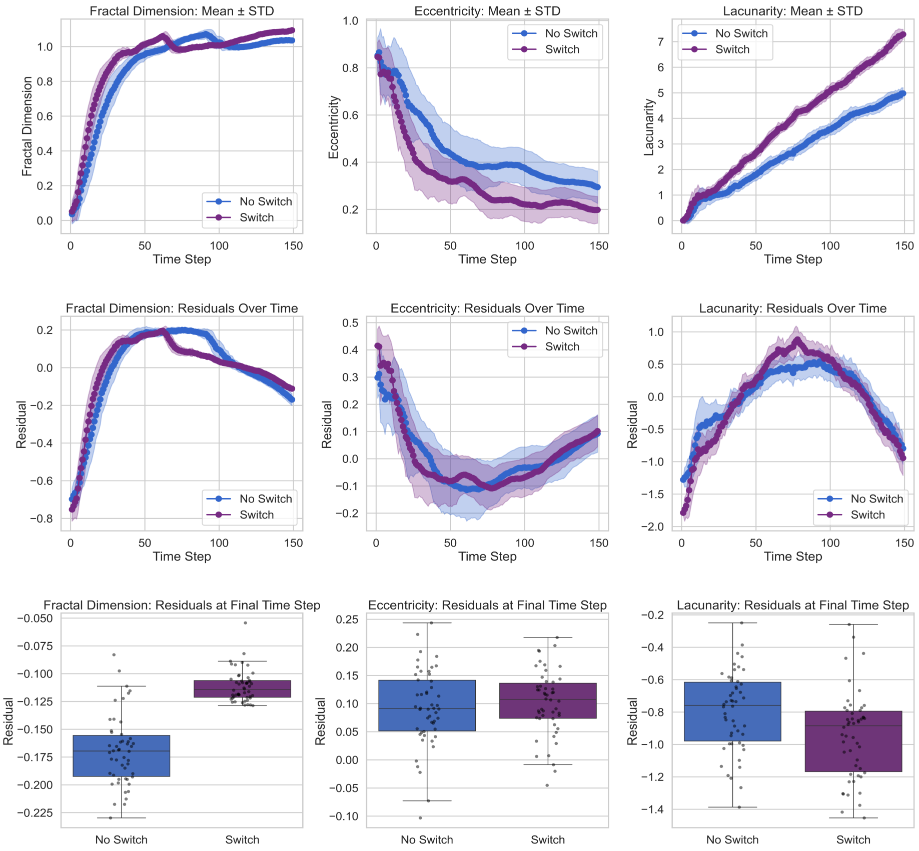


**Fig. S4 Temporal dynamics and residual morphology analysis of tumor growth under astrocyte reprogramming.** (Top row) Temporal evolution of spatial metrics—fractal dimension, eccentricity, and lacunarity—comparing model outcomes with and without astrocyte switching over 150 simulation timesteps. Shaded regions represent $\pm$ standard error across 50 replicates. (Middle row) Residualized trajectories of the same spatial metrics after regressing out tumor size at each time point, isolating morphology effects independent of tumor burden. (Bottom row) Final timestep boxplots of residual values for each spatial metric across conditions, showing that astrocyte reprogramming increases fractal dimensions, reduces lacunarity, but does not significantly alter eccentricity.

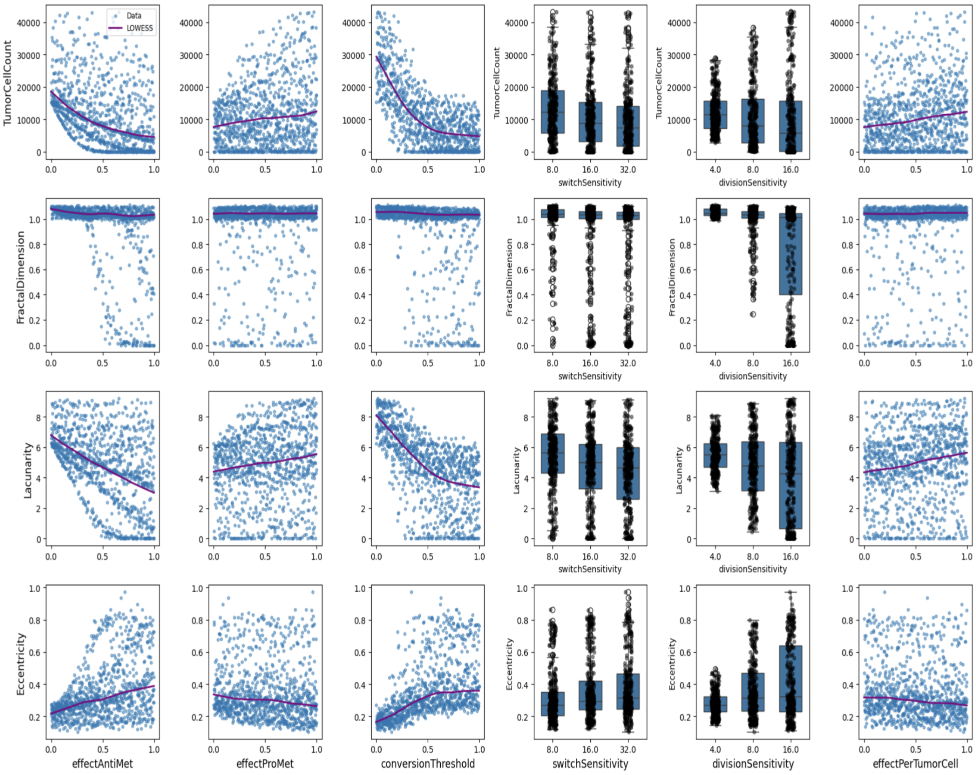


**Fig. S5 Visualization of parameter influence on tumor burden and spatial morphology metrics.** Scatterplots and boxplots showing the relationship between six key model parameters—effectAntiMet, effectProMet, conversionThreshold, effectPerTumorCell, switchSensitivity ($S_A$), and divisionSensitivity ($S_T$)—and four model outputs: tumor cell count, fractal dimension, lacunarity, and eccentricity. For continuous parameters, a locally weighted regression (LOWESS) curve is overlaid to visualize nonlinear trends. For discrete parameters ($S_A$ and $S_T$), boxplots illustrate the distribution of outcomes at each tested level. These relationships highlight how specific parameter regimes shape tumor growth and spatial complexity.

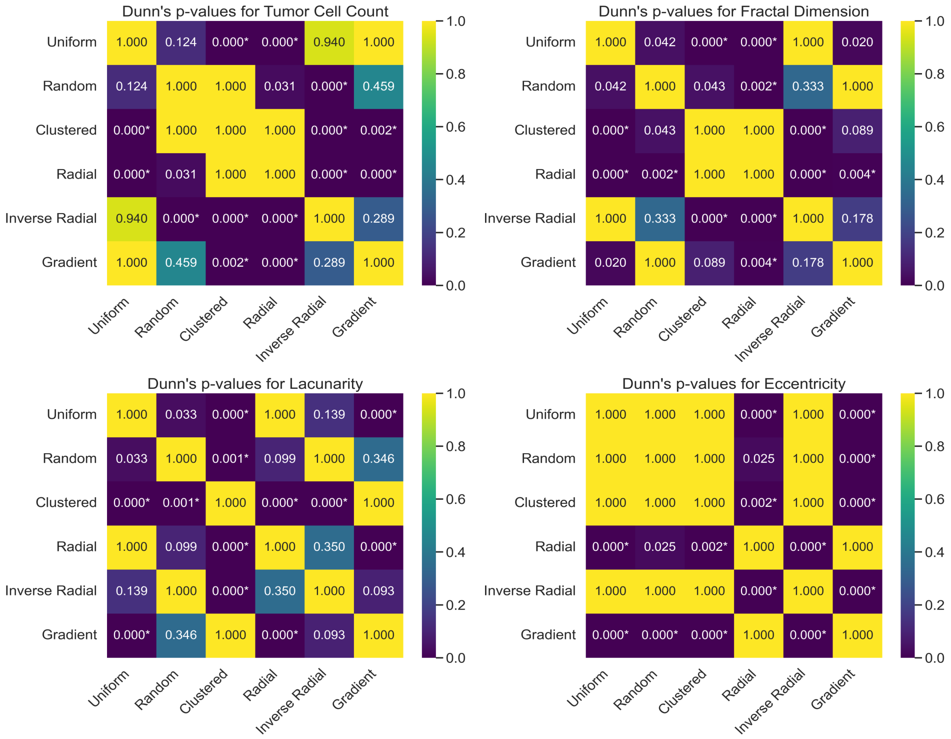


**Fig. S6 Post-hoc statistical comparisons of astrocyte spatial distributions across tumor metrics.** Heatmaps show pairwise p-values from Dunn's post-hoc test for six astrocyte spatial configurations (clustered, gradient, inverse-radial, radial, random, and uniform) across four tumor outcome metrics: tumor cell count (top left), fractal dimension (top right), lacunarity (bottom left), and eccentricity (bottom right). Lower p-values (purple) indicate statistically significant differences between spatial distributions after Kruskal–Wallis testing. Nearly all pairwise comparisons for fractal dimension and lacunarity reach significance, underscoring the strong influence of astrocyte spatial structure on tumor morphology.

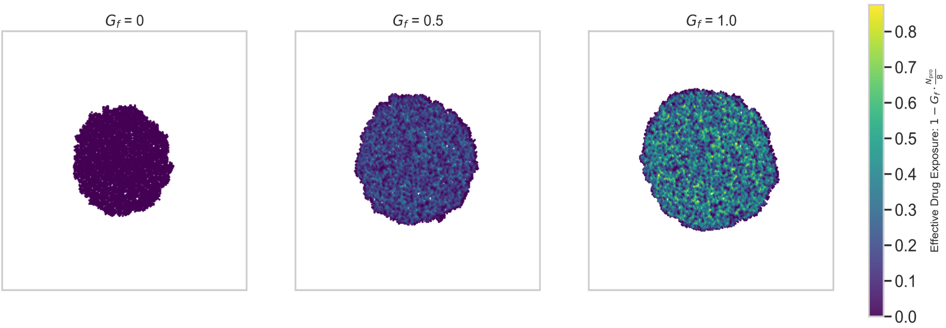


**Fig. S7 Spatial distribution of chemotherapy resistance under increasing astrocyte-mediated modulation.** Representative tumor configurations are shown for three values of the gap junction modulation factor ($G_f$ = 0.0, 0.5, and 1.0). Each tumor is colored by the level of chemotherapy resistance, computed as a function of the number of proximate pro-metastatic astrocytes. The color bar indicates resistance scaling from 0 (low resistance, purple) to 1.0 (high resistance, green). Higher $G_f$ values result in increased protection from drug exposure in astrocyte-rich regions, leading to spatially heterogeneous resistance patterns and larger tumor masses.
